# Estimate of Within Population Incremental Selection Through Branch Imbalance in Lineage Trees

**DOI:** 10.1101/002014

**Authors:** Gilad Liberman, Jennifer Benichou, Lea Tsaban, Yaakov Maman, Jacob Glanville, Yoram Louzoun

**Author notes:** Corresponding author: Yoram Louzoun, Department of Mathematics, Bar Ilan University, Hazeitim 99, Ramat-Gan 52900, Israel.

## Abstract

Incremental selection within a population, defined as a limited fitness change following a mutation, is an important aspect of many evolutionary processes and can significantly affect a large number of mutations through the genome. Strongly advantageous or deleterious mutations are detected through the fixation of mutations in the population, using the synonymous to non-synonymous mutations ratio in sequences. There are currently to precise methods to estimate incremental selection occurring over limited periods. We here provide for the first time such a detailed method and show its precision and its applicability to the genomic analysis of selection.

A special case of evolution is rapid, short term micro-evolution, where organism are under constant adaptation, occurring for example in viruses infecting a new host, B cells mutating during a germinal center reactions or mitochondria evolving within a given host.

The proposed method is a novel mixed lineage tree/sequence based method to detect within population selection as defined by the effect of mutations on the average number of offspring. Specifically, we propose to measure the log of the ratio between the number of leaves in lineage trees branches following synonymous and non-synonymous mutations.

This method does not suffer from the need of a baseline model and is practically not affected by sampling biases. In order to show the wide applicability of this method, we apply it to multiple cases of micro-evolution, and show that it can detect genes and intergenic regions using the selection rate and detect selection pressures in viral proteins and in the immune response to pathogens.

## Introduction

The phenotypic effect of genotypic changes and whether these changes affect the population dynamics still remain one of the most important questions in many domains of ecology and evolution. In a population seeded by an initial organism, mutations can affect the average number of offspring. An increase in the number of offspring is often treated as indicator for a better fitness and vice versa. Given an observed set of genes within a population, a central question arising in many domains of population dynamics is whether the observed genetic constitution of a population can be explained by a neutral random drift, or must one incorporate the effect of mutations on the fitness to explain the observed distribution of genes in the population.

This question is asked at the general level in evolution, where a debate has emerged between selection-based evolution and neutral evolution (Kimura 1968; King and Jukes 1969; Kimura and Ohta 1974). It is also often addressed at the micro-evolution level, as happens for example in viral escape mutations to avoid immune mediated destruction (Weiner et al. 1992; Allen et al. 2004; Cox et al. 2005), the dynamics of specific clones in the B cell response against pathogens (Liu et al. 1989; Berek et al. 1991) or maternal inheritance within a population (Lande and Kirkpatrick 1990; Badyaev 2005). These cases are examples of processes involving rapid asexual reproduction, where constant diversification and possibly adaptation occur with a high mutation rate.

When the effect of mutations is drastic, as is the case for strongly deleterious or advantageous mutations, a clear genetic signature of the selection can be observed in the genome, and multiple methods have been proposed for measuring selection in such cases. Some of these measures rely on the ratio of synonymous (S) to non-synonymous (NS) mutations. Specifically, a comparison of the observed and expected NS/(NS + S) ratios is often used as a measure for selection. The expected ratio is calculated based on an underlying mutation probability model (e.g. (Nei and Gojobori 1986; Yang 1998; Yang and Nielsen 2000)), or based on genetic regions where no selection is assumed to occur (Shlomchik et al. 1987). An increased frequency of NS mutations is treated as an indication for positive selection and vice versa. These methods are often useful, when a good estimate of the baseline mutation model is available. They may however lead to erroneous conclusions when the baseline mutation model (i.e. the expected probability of each mutation type) is inaccurate, as happens for example in immunoglobulin sequences (Hershberg et al. 2008).

In many cases of micro-evolution, the observed time scale of the dynamics is limited, and the fitness (dis)advantage induced by mutations may be limited. In such a case the fixation probability will be low, and S to NS based methods will be less useful. A different approach proposed for detecting selection in such cases is to use properties of lineage trees. Two of the most powerful such measures proposed for the detection of selection (Maia et al. 2004; Li and Wiehe 2013). are Sackin’s and Colless’s statistics (Sackin 1972; Colless 1982; Kirkpatrick and Slatkin 1993; Blum and François 2005). Sackin’s index is the average root-leaf distance (over all leaves). Colless’s index is the sum of imbalance over all nodes, where a node’s imbalance is taken to be the difference in number of leaves between the bigger and smaller sub-trees. These measures are tested vs. a neutral model, which is usually the Yule model, where a tree is constructed by giving each branch the same probability to split (Yule, 1925). Other statistics do not use trees but are based the number of segregating sites, most notably Tajima’s D (Tajima, 1989).

These methods have two well-known limitations. They do not distinguish between S and NS mutations and statistical power is lost. Most of these methods measure deviation from a neutral model and cannot differ between different types of selection, e.g. positive and negative ones.

We here offer a more direct approach to measure incremental selection within a specie passing a continuous adaptation, which is directly related to a quantitative definition of the meaning of incremental selection. This new method overcomes limitations of the S to NS mutation ratio and of the tree shape based selection detection methods, by accounting for the completing information found in each of the two, that is, the classification into mutation types, and the imbalance between different sub-trees.

## Results

### Selection

Assume a population originating from a single founder through division, with a given ancestral sequence in the region of interest (a gene, a combination of genes or even a part of a gene). Mutations in this region can potentially affect the population dynamics. In such a case we would define positive selection in the population as an increased average division/birth rate or a decreased average death rate following mutations (note that these are not precisely the same (ANDERSON *et al*. 2009), but the distinction is beyond the scope of the current analysis). A decrease in the division rate would be defined as negative selection. Obviously, each mutation by itself can have a positive, null or negative effect, but the definition of selection is based on the average population dynamics and not with the dynamics following a single mutation.

Let us follow a mutation that occurs within a population, if this mutation increase the average number of offspring per generation from *μ* to *μ* + Δ*μ*, then by a time proportional to log of the total population size, the advantageous mutation will take over the population (KIMURA 1962), and when we will compare the population to its latest common ancestor (LCA), we will have no direct evidence that such a mutation has occurred (Fig 1A,B). In this case the genetic composition of the population would be equivalent to the one expected in a neutral model. The only difference would be the addition of a single NS mutation to a gene in the entire population. If external reference (e.g. from the comparison to orthologues) that can help us define the sequence seeding the population are available, they can be used to infer that selection has taken place.

**Figure 1.**
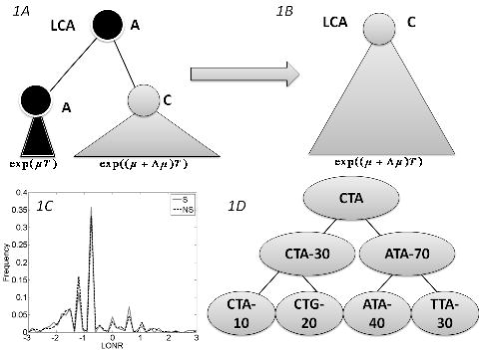
The branch imbalance framework and examples. (A) Schematic view of a branch corresponding to a mutation event. Following a mutation, the population can be expanded (or reduced), the advantage will lead to an exponentially growing difference in the number of offspring in parallel branches descending from the same origin (B) After some time, one branch will take over the entire sample, and the information carried in the ratio between the branches will be lost. (C)LONR values histogram for one simulated sequence pool, simulated under naive multiplication from unique ancestral sequence. While the average is not 0, there is no difference between branches following S and NS mutations. (c) LONR values histogram taken from mouse data (see main text for further details) exhibiting positive selection for R mutations. (D) Example of tree. In the left branch a mutation occurred from CTA to CTG, and the ratio between the mutated and un-mutated branches number of offspring is 20/10. In the right branch, a mutation from ATA to TTA occurred, with a ratio of 30/40. In the root, a mutation from CTA to ATA occurred with a ratio of 70/30.

However, in many cases, evolution occurs over an intermediate period and is weak, leading to the coexistence of the two alleles in the genome of the population (the mutated and the un-mutated one).In such cases, we expect the ratio between the two allele frequencies to be proportional to *e*^Δ*μT*^, where T is the time from the mutation to the sampling time (Fig. 1A).

For a single mutation, it is impossible to differentiate between the effect of selection and a non-uniform sampling where one branch is sampled more deeply. However, if many mutations occur in the genetic region of interest, and if, in average, mutations in this region increase the average number of offspring, we expect, in average, more offspring in branches that follow a mutation in this region than in branches emerging from the same direct ancestor with no mutations, and inversely in the case of negative selection.

We thus propose to detect incremental selection using this imbalance in cases where most mutations are neither strongly deleterious nor strongly advantageous and where the time scale studied is too short to allow the fixation of slightly advantageous mutations. Such cases are far from being rare and become more and more frequent as the depth of genetic sampling increases in many domains (Metzker 2009; Kircher and Kelso 2010; Ozsolak and Milos 2010; Nielsen et al. 2011; Benichou et al. 2012).

### Effect of sampling

Assume a sample from a population dynamics process, with a “real lineage tree” representing the actual division and mutation process. In the real tree, the average ratio between the number of leaves under an internal node that has a given mutation and the parallel descendent of their common direct ancestor that does not have a mutation (i.e. its un-mutated sibling) should be 1. The same cannot be told of the reconstructed lineage tree based on the sampled distribution, following biases induced by the sampling or the tree construction algorithm (Fig. 1C). More specifically, a branch with a specific mutation is one possible offspring out of many. Thus, this specific branch will be typically smaller than the parallel branch holding the other offspring. However, in the absence of selection, the ratio between the total number of offspring of a branch with a mutation and without one should be similar following S and NS mutations. Thus, the in order to estimate the presence of selection, one can simply compare this ratio (that we denote as the branch size imbalance) following S and NS mutations.

### LONR

We define a measure of selection in a gene as the ratio of the number of leaves (measured descendants) under a branch where a mutation occurred and the number of decedents in its direct sibling where no such mutation occurred. We compare the distribution of these ratios (more precisely the log of the ratios) in all S and NS mutations to estimate whether the distribution deviates from the one induced by neutral drift (Fig. 1D).

Specifically, for each mutation occurring in one son of an internal node and not in the other, we compute the sub-tree size under the son with a mutation and the sub tree under the son without a mutation. Positions where a mutation occurred in the two sons (e.g. A->C and A->G) are ignored. The log of the ratio between these two sizes is defined as the Log Offspring Number Ratio (LONR) of this mutation. We then compute the LONR value for all S and NS mutations in the tree, and compare the S and NS LONR distributions (Fig S1, S2). These mutations are computed on the reproduced lineage tree, which may differ from the real tree. The effect of the tree production method will be further discussed.

The mutations of interest can be all the mutations occurring in a gene, a gene combination, or even a genetic region composing a part of a gene that can be continuous or discontinuous. Formally, we define a set of positions in a genetic segment, and only count mutations in this region (Fig S1, S2).

Note that this analysis is not sensitive to the details of the baseline model for the probability of either S or NS mutations, since their absolute number is never used in the analysis. The only case where such a model would affect the current measurement is in the extreme case that the baseline mutation probability would induce a much higher S than NS probability

### Simulated data

In order to check that the LONR does not detect selection in its absence, we simulated a Yule process, sampled the resulting sequences (see Methods for details), produced lineage trees and compared the LONR distribution following S and NS mutations (Fig 2A). In the regime of over 10-20 mutations per sequence and at least 300 sequences per tree, the False Positive (FP) rates (the cases where the LONR average is significantly different following S and NS mutations with a p-value of 0.05) are near the expected 5% (Fig. 2A). This range of mutation and sequence numbers is typical to most current applications of lineage trees and phylogenetics. We here limit the analysis to this range.

**Figure 2.**
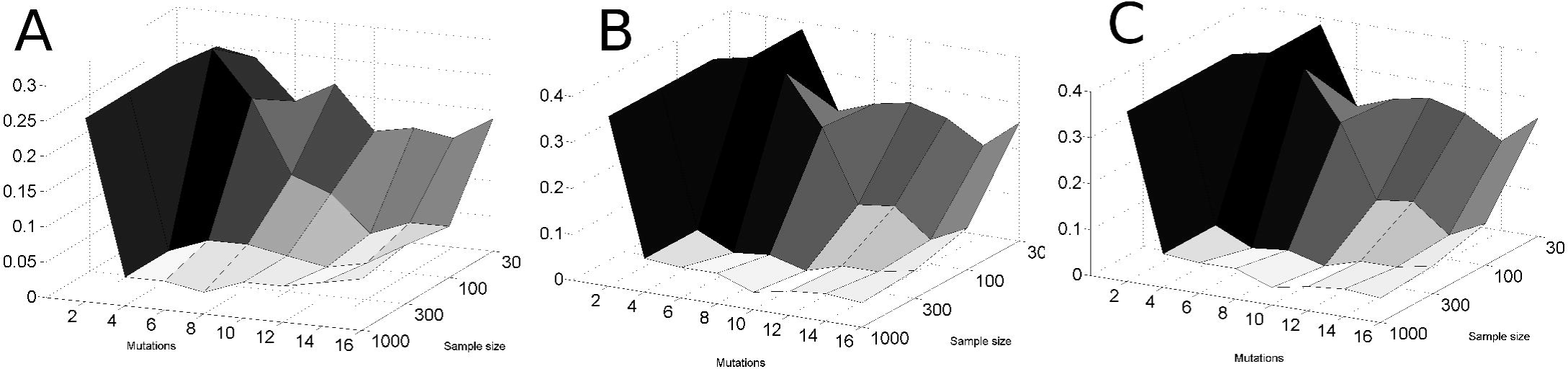
Fraction of lineage tree where selection was detected at a p = 0.05 level, as a function of the average number of mutations per sequence and the sample size. In all cases, the false positive fraction is around 0.05 (as expected randomly), when the sample size is above 300, and when there are at least 4-5 mutations per sequence. The results are consistent for a uniform mutation rate (along the sequence) (A drawing), non uniform mutation rate, with some regions having a twice higher mutation rate (B drawing), as well as when the mutation rate is non-uniform and the sampling is non uniform (C Drawing).

We have repeated the analysis with non-uniform mutation rates (position dependent mutation rates) and with sampling biases, and obtained similar results, as long as the S and NS mutation rates are of the same order of magnitude (Fig 2B and 2C). Specifically, sampling bias was simulated by oversampling descendants from one of the clones of the 3rd generation. Again, in the domain of 300 sequences or more and an average of 10 mutations per sequence or more, the sampling effect and the non uniform mutation rates along the sequence did not increase the error rate (See methods for mutation and sampling models).

We avoid a major sampling bias by averaging over mutation events and not over sequences. Suppose for example, we would analyze a clone that led to two populations, one much more sampled than the other. This would affect a single internal node (probably the root), but the imbalance in all the other nodes would be unaffected. Within this internal node, the effect of over-sampling would be similar in S and NS mutations. Sub-sampling would have a significant effect only through the combined second-order effect of the sub-sampling in one node combined with the difference in the S and NS mutation frequencies. This effect is of no practical importance in all the examples studied here and probably in most realistic situations.

### Mitochondrial sequences

A typical case where the number of generations is low and the mutation rate is high is maternal inheritance in the human mitochondrial genome. We sampled 3,106 sequences from published mitochondrial genomes in the NCBI database that passed our quality validation checks (see Methods). We computed the average LONR value over all positions using a sliding window of 400 nucleotides and 95% overlap. When looking at the LONR score for all mutations (both S and NS), the distribution is non-uniform with very large peaks in the total LONR score. These peaks overlap with the known mitochondrial genes as well as an rRNA region in positions 1671-3229 (Fig. 3 grey line). In other words, the LONR delineates important regions in the mitochondrial genomes, where mutations have an important effect, with no a-priori knowledge. Specifically, a strong positive selection force is present in the area between 1671-3229 nucleotides, which codes for the 16S ribosomal RNA that has been suggested to undergo strong adaptive selection for mutations affecting stem-loop secondary structure of the ribosome (Ruiz-Pesini and Wallace 2006).

**Figure 3.**
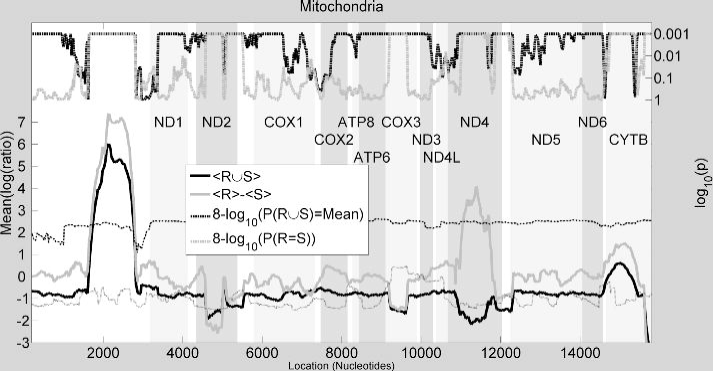
Mutation and selection pressure in full mitochondrial genomes. Mean LONR values for (solid grey line) all mutation events along with (dashed grey line) p-value of one-sample t-test for divergence from the overall mean, and (solid black line) difference in mean LONR values for NS and S mutation events along with (dashed black line) p-value of two-sample t-test for difference between NS and S mutation events. The data was processed using sliding-window scheme with bin size of 400 nucleotides and 95% overlap. The highlighted bars are the mitochondrial genes. One can clearly see that the bands of selection follow closely the positions of some of the genes. The selection bands are narrower than the genes, following the effect of the sliding window. The drop at the end is a boundary effect. The dashed dotted and dotted thin lines are the Tajima’s D index and the NS/(NS + S)-NS0/(NS0 + S0) index. The two indices do not detect selection in the ribosomal RNA and are not sensitive to the precise positions of genes for most genes.

In gene regions, the S and NS LONR scores were compared using a t-test and an FDR correction (Benjamini and Hochberg 1995) was applied. In most genes and in the rRNA, the difference from the baseline is significant (p < 0.001 t-test) (dark full and dashed lines in Fig. 3). Among the 13 coding regions, there are some prominent areas such as CytB, ND4 where positive selection takes place, and ND2 and COX3 that undergoes negative selection. In the ribosomal RNA, we do not compute an NS to S difference, since we cannot clearly define NS and S mutations. Applying either the Tajima’s D index or an S to NS measure on the same sequences does not clearly provide a distinction between genes, and does not detect the Ribosomal RNA (Fig 3. dotted and dashed dotted thin lines).

Previous studies have measured selection in mitochondrial genes. The methods by which selection was detected in these studies were either S to NS mutation ratio (Kivisild et al. 2006), relative selective constraint (Mishmar et al. 2003; Kivisild et al. 2006) or neutrality index (Elson et al. 2004). Our results agree with most of the literature. Measures for selection on CytB and Cox3 were consistent with our observation: CytB was consistently found to undergo positive selection (Mishmar et al. 2003; Kivisild et al. 2006; Ingman and Gyllensten 2007). Similarly, COX3 was shown to have relatively low S/NS ratio and high neutrality index (Mishmar et al. 2003; Elson et al. 2004; Kivisild et al. 2006; Ingman and Gyllensten 2007) suggesting a negative selection on this gene. Moreover, for most genes where we did not discover selection, no stringent selection was reported in the literature. Similarly, as described by (Ruiz-Pesini and Wallace 2006), mutations are systematically positively selected in the ribosomal RNA. Note that the LONR can provide a very clear estimate of the strength of selection in this region, and it is much stronger than in regular genes.

Still some differences exist between some published results and the LONR measure. Mainly that ND4 is claimed to undergo negative selection, and ND2 to undergo positive selection in contrast with our study. ATP6 is reported to pass selection, which is not detected by the in LONR measurement. The source of the difference is probably that we measure relative selection within a population, while NS/S measures are affected by genes and alleles common to the entire population. In other words, traditional measurements estimate whether mutations increase the probability of a cell with a sequence carrying this mutation of being observed, while we measure whether mutations increase the growth rate a sub-population carrying it. At the tree measurement level, we are interested in the effect of replacements in respect to direct father, and not in the difference between a sequence and a remote ancestral sequence.

### Viral sequences

Another interesting case of population dynamics with a high mutation rate, and an expected strong selection is the escape of viruses from detection by the immune system through mutations in their epitopes. RNA Viruses accumulate mutations at a rate of one mutation per division per genome approximately. We analyzed 21 viral proteins from 4 organisms (see Methods), and computed for each protein the difference between the S and NS LONR distributions (Table 1, Fig. 4). Proteins were divided into CD8+ T cell epitope and non-epitope regions (see Methods for a detailed explanation of epitope description).

**Figure 4.**
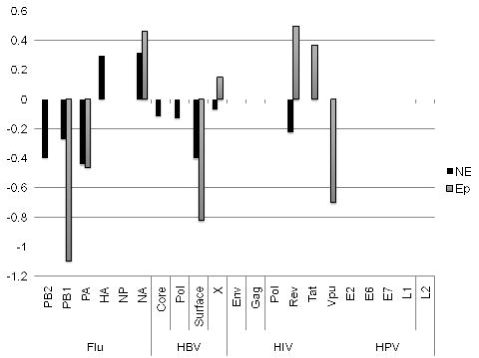
Difference between LONR score for NS and S mutations inside (Ep) and outside (NE) T cell epitopes. Only cases with a t-test p value of less than 0.05 are drawn. All values are given in Table 1. The positive values represent positive selection, and negative values represent negative selection. Positive selection is observed in proteins known to void recognition by the immune system.

**Table 1.** Mean LONR NS-S differences for multiple viral proteins, separated into epitope (Ep) and non epitope (NE) regions. The highlighted scores have a p value of less than 0.05. As can be clearly seen, the selection is negative in most proteins and most regions is negative, as expected if viruses have reached an optimal sequence a long time ago. However there are some regions of positive selection, especially in regions where the immune response drives the accumulation of escape mutations.

|  |  | NE | Ep |
| --- | --- | --- | --- |
| Flu | PB2 | **-0.39859** | 0.122483 |
| Flu | PB1 | **-0.27061** | **-1.09592** |
| Flu | PA | **-0.4446** | **-0.46262** |
| Flu | HA | **0.292482** | 0.010707 |
| Flu | NP | -0.04631 | 0.264885 |
| Flu | NA | **0.315675** | **0.461664** |
| HBV | Core | **-0.11408** | -0.18042 |
| HBV | Pol | **-0.12908** | -0.08908 |
| HBV | Surface | **-0.40382** | **-0.82322** |
| HBV | X | **-0.06924** | **0.147685** |
| HIV | Env | 0.01211 | -0.10057 |
| HIV | Gag | 0.029754 | -0.04905 |
| HIV | Pol | 0.032609 | -0.08632 |
| HIV | Rev | **-0.22601** | **0.496061** |
| HIV | Tat | -0.09679 | **0.36844** |
| HIV | Vpu | -0.20211 | **-0.70019** |
| HPV | E2 | 0.004034 | -0.03717 |
| HPV | E6 | 0.020337 | -0.2272 |
| HPV | E7 | -0.12331 | -0.01 |
| HPV | L1 | -0.09061 | 0.038934 |
| HPV | L2 | -0.02724 | -0.15036 |

A strong and significant negative selection was observed in multiple proteins, both in epitope and non-epitope regions (Fig 4) in HIV, Flu and HBV, but not in HPV. Such a selection is expected if viral proteins have reached an optimal sequence a long time ago.

In some epitopes, a clear positive selection has been observed in the epitopes (t-test p-value <0.001 for HIV TAT and HIV Rev) in the two main proteins reported to mutate away their epitopes (Addo et al. 2002; Betts et al. 2002; Kiepiela et al. 2004; Vider-Shalit et al. 2009a). Note that this observed selection represents the rapid removal of epitopes and not the removal of epitopes that may have occurred historically, since we only look at mutations that can be computed from the current sequences compared with their LCA. Outside T cell epitopes, a positive selection is only observed in the Influenza Hemagglutinin and Neuraminidase, which are known to accumulate mutations to avoid the detection by antibodies. Thus the LONR indeed detects the best known targets of positive selection in viruses. Note that it does not detect an advantage for escape mutations in other proteins. There, this advantage may be too weak, or masked by a parallel negative selection, yielding a net unobservable detection.

When using either the Tajima’s D index or the NS/S measure, systematically, more positive and negative selection are observed outside epitopes than inside epitope (t-test on the absolute value of D or [NS/NS+S]-[NS0/NS0+S0] between epitope and non-epitope regions p<0.001, Fig S3). Moreover, negative selection is observed in practically all viral genes, when using the NS/S method (t-test of all viral proteins vs. 0, p<1.e-4, Fig S3). This is probably due to an inaccurate baseline model.

### Mouse Immunoglobulin

Probably the most classical real time evolution with a high mutation rate and growth of clones is the affinity maturation process of B cells in germinal centers. In this process, an initial B cell grows into a clone and during its growth hyper-mutations occur in the B cell receptor at a rate of one mutation per division (Kleinstein et al. 2003) and an extreme division rate (Anderson et al. 2009). They thus fit precisely the LONR framework. We have studied two mice strains: one starting with a high affinity to the experimental antigen tested, and one with a low affinity. In order to induce a potent immune response, the low affinity mouse strain must accumulate a large number of specific mutations to obtain a high enough affinity to its receptor (Dal Porto et al. 2002). The mouse strain with an initially high affinity can form clones even with the germline receptor it has, and it thus intuitively not under a very stringent selection.

Indeed, the initially low affinity mice show a large difference between S and NS mutation LONR scores, with a clear positive selection, while the high affinity mice do not show such a difference (Table 2). Note that in principle these mice could also have a negative selection, where mutations in average reduce the fitness of the cells. However, we have not observed such a selection in these mice strains. While S/NS methods have been used in Immunoglobulin data (Nei and Gojobori 1986; Nemazee 2000; Kleinstein et al. 2003; Anderson et al. 2009), their results have been shown to be very sensitive to complex baseline model of Ig mutations, and errors in the model led to many erroneous conclusions on the presence or absence of selection (Hershberg et al. 2008).

**Table 2.** Mean LONR NS-S differences and two-sample t-test values for the two mouse types, calculated using four tree construction algorithms. The results are similar for most algorithms, except for the UPGMA, which is quite simplistic and often contains unrealistic assumptions, such as a uniform molecular clock.

|  | R-S LONR |  | p-value |  |
| --- | --- | --- | --- | --- |
|  | Difference |  |  |  |
|  | Lyle | Meg | Lyle | Meg |
| Maximum parsimony | 0.6243 | 0.1926 | 0.0095 | 0.2766 |
| Maximum likelihood | 0.7475 | 0.1895 | 0.0141 | 0.4502 |
| Neighbor joining | 0.5174 | 0.1945 | 0.1113 | 0.5154 |
| UPGMA | -0.3524 | -0.0267 | 0.2813 | 0.9294 |

### Human Immunoglobulin

A more interesting case is the full B cell repertoire of a human host. In such a repertoire two opposite forces operate: a) mutations can ruin the functionality of the receptor and decrease its survival probability, and b) mutations can on the other hand increase the affinity to the antigen and thus lead to a higher division rate. The Complementarity Determining Region (CDR) of the B cell receptor determines its interaction with the antigen, and mutations there have a higher probability to increase the affinity than mutations in the framework (FWR) region (Berek et al. 1991; Cowell et al. 1999). However, the net selection effect in each of these regions still remains unclear. Beyond the effect of somatic hyper-mutation, B cells are affected by isotype switches from naïve IgM to memory IgM, and from there to memory IgG and IgA. The memory (IgM, IgG and IgA) isotypes occur at the advanced stages of the immune response and thus lineage trees based on such receptors are expected to represent the full evolution following selection.

We have used high-throughput sequencing to sequence over 500,000 B cell receptor samples from each donor, in 12 donors. We built lineage trees from the sequences (See (Benichou et al. 2013) for details of sequences, and production of lineage trees), and measured the LONR distribution in all IgA and IgG sequences trees and compared the LONR distribution in NS and S mutations. At the first stage, we only analyzed trees with significantly different NS and S LONR averages (unpaired two-sided t-tests, p < 0.01), and analyzed two regions of the B cell receptor where the junctional diversity had no effect on the construction of the lineage trees: FWR3 and CDR2 (Lefranc et al. 2009). The results are quite striking. As expected in both IgG and IgA memory cells, the positive selection is much stronger for the CDR region than for the framework (Fig. 5). However, even the FWR region passes a positive selection during the immune response. Such a positive selection in the FWR region suggests that the large clones (i.e. clones that were selected to grow more than others), are actually affected by structural changes in the FWR region. This selection may represent the need for structural changes in the immunoglobulin structure to reach a very high affinity.

**Figure. 5.**
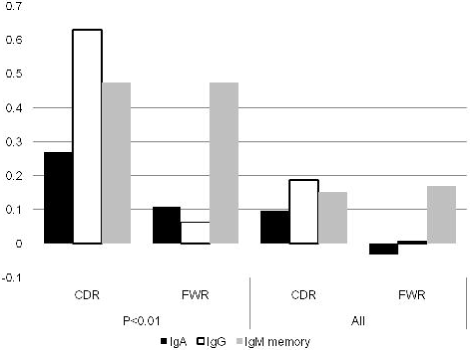
Mean LONR values for immunoglobulin sequence pools. Mean LONR values for memory IgM, IgA and IgG sequence pools, where the values are averaged over (a) all trees in which difference between overall NS and S LONR mean values was found to be significant (t-test, p < 0.01) and (b) all trees.

Interestingly, when analyzing all trees (over 30,000 lineage trees), the reported negative selection in the FWR region (Hershberg et al. 2008) appears in the IgA isotypes (Fig 5). This leads to the interesting conclusion that selection may be affecting differently the main part of the distribution and its extremities. In the main part of the distribution, non-synonymous mutations in the CDR are selected, since they improve the affinity, and non-synonymous mutations in the FWR are selected against since they ruin the structure of the antibody. In the extreme cases, the non-synonymous mutations in the FWR are also selected, since some of these mutations can actually improve the affinity and enlarge the resulting clones. A much more detailed analysis of this specific dataset can be found in (Liberman et al. 2013). The Ig analysis is a classical example of the simultaneous positive and negative selection in different regions of the same gene and of the possibility of detecting such selection using the LONR.

### Effect of tree building algorithm

Constructed lineage trees are only estimates of the real lineage, and their precise shape may be sensitive to the algorithm used to build them as well as to the baseline mutation model. We have thus tested whether the methodology used to build the trees affects the LONR scores. We have constructed the lineage trees from the two mouse strains discussed previously using four methods: Maximum Likelihood, Maximum Parsimony, Neighbor Joining and UPGMA. All algorithms were applied using the Phylip toolbox (Felsenstein 1989). In all methods, except for UPGMA, the LONR results were similar, with the maximal difference between S and NS mutations being in the MP algorithm. UPGMA is a highly simplistic algorithm and should not be used to detect fine details of tree shapes.

## Materials and Methods

### Alignment and Phylogenetic trees

The DNA sequences of different viruses were aligned using the TranslatorX program (Abascal et al. 2010) that aligns nucleotide sequences based on their corresponding amino acid translations. Phylogenetic trees were then produced from the aligned sequences using the maximum parsimony method of the Phylip bioinformatics tool package (version 3.69) (Felsenstein 1989). For the mice data, three other tree construction techniques, neighbor joining, maximum likelihood and UPGMA, were used to validate robustness to construction algorithm. For samples with over 100 sequences, a neighbor joining algorithm was used in the same package. For each group of sequences, a genetically distant ‘outgroup’ sequence was added to position the root of the tree, and reconstruct the ancestral sequences. To avoid ambiguous nucleotides in internal nodes, when both child sequences had a gap in a certain locus, the parental nucleotide was changed to a gap as well. If one of the child sequences had a non-ambiguous nucleotide, the parental nucleotide was changed accordingly.

While recombination may be important in general in viruses (Wilson et al. 2009), we have ignored its effect, and did not find evidences for it in our current dataset. In the mitochondrial dataset and the Ig datasets, a single lineage tree was built for each group of sequences. The separation into regions was performed after the construction of the lineage tree.

### Selection Score

Given a tree, each mutation event is assigned: (a) a NS or a S mutation flag by its effect on the amino-acid translation of the containing codon; (b) the location of the mutation (related gene where applicable, and number of nucleotides from the beginning of the sequence, otherwise); and (c) The log of the ratio between the number of leaves (sequences) in the sub-tree following the mutation branch and the number of leaves in the sub-tree following the non-mutated branch (see Fig. 1, and Figs. S1 and S2). This ratio is denoted the Log Offspring Number Ratio (LONR). This log-ratio is thus positive if the number of final sequences marked by the tree construction algorithm as descendants of the mutated sequence is larger than the number of final sequences marked as descendants of the non-mutated sequence, suggesting some better fitness of the mutated sequence, or positive selection for such mutation, and negative in the opposite case. For each area of the sequence, a t-test is performed (unpaired, unequal variances) between the NS and S mutations.

### Simulation

A sequence pool simulating neutral reproduction was generated from a random original sequence of 348 nucleotides, with a constant multiplication rate of two offspring per organism. Two equal size regions (174 Nt. each) were defined with uniform mutation probabilities with average mutation rate of 1/2 and 1 mutation per generation. The population was sampled in different sample sizes and along different generations. In each sampling, one of the eight first siblings (the third generation) was chosen randomly, and its descendants had a twice higher probability of being sampled, effectively simulating sampling bias for a specific clone. The process was repeated 1,000 times, and selection was computed in the described process. NS and S mutations were defined relative to their direct ancestor, resulting in unequal NS and S probabilities. All mutations had equal probabilities (i.e. we did not make a difference between Purines and Pyrimidines). We tracked all sequences in the simulation, and the last generation of the simulation was sampled to produce the lineage trees.

### Statistical Analysis

For the mitochondrial sequences, the analysis was performed using a sliding window of 400 nucleotides and 95% overlap. The p-values are presented for each window, along with the differences in the mean LONR values between the NS and S mutations. In order to asses areas where selection forces are presented for NS and S events alike, a one sample t-test is performed on the whole mutation events LONR values, for NS and S together. When reporting the final results, a FDR correction was performed to account for the large number of windows (800).

For the viral sequences, a single tree was constructed for each virus from the obtained sequences, and a two-way ANOVA test was performed for assessing the significance of the NS vs. S grouping, the epitopes vs. non-epitopes grouping and interactions between the two.

For the transgenic mice data, trees were constructed for different clones and LONR values were collected from all trees, grouped by the two mouse types. Mean NS-S LONR values are reported along with two-sample t-test p-values.

For the immunoglobulin data, the receptors where clustered by isotype (IgA and IgG). Lineage trees were constructed and the sequences were divided to CDR and FWR regions. Mean LONR NS-S difference was computed per clone and per region along with two sample t-test p-values.

### Viral and mitochondrial Sequences

All sequences were obtained from the NCBI nucleotide database (Benson et al. 2004). We have used sequences from Influenza A (1,000 sequences for segment 1 to segment 6), HBV (1694,2370,211 and 999 sequences for Core, Polymerase, Surface and X, accordingly), HIV (179,823,731,159,757 and 150 for Env,Gag,Pol,Rev,Tat and Vpu accordingly), and HPV (105, 89,88,72, and 121 for E2,E6,E7,L1 and L2, accordingly). For the sake of lineage trees design (see the next section), we have defined an outgroup for each set using genetically distant homologues (e.g. Influenza B for Influenza trees).

The human mitochondrial sequences were all of the full genome nucleotide sequences available at the NCBI, with a length of at least 15574 and at most 16581 nucleotides. 2689 sequences were used with hosts from multiple regions including large cohorts from China and India.

### Mouse Data

The sequences from transgenic mice were obtained from two H chain transgenic mice (Hannum et al. 2000; Dal Porto et al. 2002) that were backcrossed with Jh KO/Balb mice (Chen et al. 1993; Hannum et al. 2000) for nine or more generations. All mice were maintained under specific pathogen-free conditions and sacrificed at 6-10 week of age. Mice were immunized i.p. with 50μg of NP25-chicken gamma-globulin (CGG) precipitated in alum or precipitated alum alone as a control. B cells were sequenced from microdissections in germinal centers of these mice, 16 days after the immunization. One mice type had an initial low affinity for the antigen, while the other had an initial high affinity.

### Immunoglobulin sequences

Over 500,000 B cell receptors were sampled from each donor in 12 donors (Benichou et al. 2013), using 454 sequencing and a RACE protocol. The details of the sequencing and the validity checks are beyond the scope of this manuscript. For each sequence, the most fitting V, J, and V-J distance was found by maximizing the relative number of non-mutations for both V and J segments. Only sequences that matched higher than 0.5 in both segments were kept for further analysis. The sequences were then clustered according to the most fitting V and J as well as the distance between V and J, and were truncated to 159 nucleotides from the end of the germline V and 20 nucleotides from the beginning of the germline J.

### Defining epitope regions

Epitopes were computed using three algorithms: a proteasomal cleavage algorithm (Ginodi et al. 2008), a transporter associated with antigen processing (TAP) binding algorithm (Peters et al. 2003), and the MLVO major histocompatibility complex (MHC) binding algorithm (Vider-Shalit and Louzoun 2010). We have computed epitopes for the 39 most common human leukocyte antigen (HLA) alleles and weighted the results according to the allele frequency in the global human population. The algorithms’ quality was systematically validated vs. epitope databases and was found to induce low false positive (FP) and false negative (FN) error rates. These algorithms were validated in multiple previous analyses (Louzoun et al. 2006; Vider-Shalit et al. 2007; Almani et al. 2009; Vider-Shalit et al. 2009a; Vider-Shalit et al. 2009b; Kovjazin et al. 2011; Maman et al. 2011a; Maman et al. 2011b).

Each ninemer, in each aligned sequence, was scored according to the weighted frequency of alleles to which it binds. For longer sequences, each position in the sequence was scored according to the maximal score given to any ninemer containing it. For the whole aligned sequence population, these values were averaged on a per sequence manner, resulting in an epitope score per position. The positions that scored in the higher 15% were defined to be epitope related areas.

## Discussion

The detection of selection is a crucial issue in population biology, evolution theory and ecology. It also has important clinical implications. While multiple sequence based methods have been proposed to detect selection (Yang and Bielawski 2000; Plotkin et al. 2004; Wong and Nielsen 2004; Massingham and Goldman 2005; Pond and Frost 2005; Zhang et al. 2005; Hershberg et al. 2008), most of them are focused on strongly advantageous or deleterious mutations. We have here proposed a method best adapted to the detection of slightly advantageous or deleterious mutations in micro-evolution.

The basic concept behind the here reported LONR measure is to test for the systematic increase of the population size following non-synonymous mutations in a given regions. An advantage of the proposed method is that each mutation is counted once independently of the total number of sequences that end up containing this mutation. Thus, it is practically not affected by sampling biases or by the expansion of specific sub-populations.

While multiple tree shape based methods were developed (Sackin 1972; Colless 1982; Kirkpatrick and Slatkin 1993; Blum and François 2005), these methods often cannot detect the direction of selection, and cannot detect which region in the sequence is selected. Moreover, many of these tree shapes are sensitive to sampling effects making them impractical to use in realistic situations (Stam 2002).

We have here proposed a new method that can clearly detect positive and negative selection or their combination, based on the effect each mutation has on the number of offspring in the tree under the branch where the mutation has occurred. This method can only be applied where the mutation rate is high enough for alleles with disadvantageous mutations to exist in the population. In other words, the mutation rate should be higher than one over the log of the sampled population size. Such a range exists for example in the population dynamics of mitochondria within a host specie, in viral dynamics and in the affinity maturation process in germinal centers. We have here studied all these cases and have shown that indeed selection can be detected in all cases studied. Other applications of this method can be the evolution of the Y chromosome and the changes in Short Tandem Repeats (STR) frequencies in it or the evolution of bacteria in an infection in the population.

We have validated that this method is precise in the domain of a large number of mutations per sequence (>10) and large samples (>300). In this domain, the method proved to have many important applications, such as the detection of selection in genes (and actually the direct detection of genes), the detection of viral proteins passing positive and negative selection and understanding the selection process in a B cell immune response. We have shown that while the ribosomal RNA has a very strong positive selection, some genes pass positive selection, and others negative selection. In B cells, we have shown that while CDR mutations are always selected, FWR mutations are selected against in the majority of the populations, but actually strongly positively selected in the extreme cases.

The comparison between S and NS mutations is only the most basic distinction. Other possibilities exist, especially change/no-change of some amino-acid property, such as size or hydrophobicity. Such methods would test for selection for specific changes and not selection for mutation in general. In other words, the proposed methodology can be used to estimate whether changes in a given property increase or decrease the number of offspring, compared with a random change.

The main limitation of the current score is that is blind to strong selection. Once a mutation is fixed in (or completely removed from) the population, we will not observe the polymorphism at this site that allows us to compare the branch sizes. This may actually be the case in the long term evolution of advanced creatures, where we do not observe inter-species.

